# Proteomic and transcriptomic host biomarkers for detection of pleural tuberculosis

**DOI:** 10.1101/2024.10.08.617199

**Authors:** Matheus Rogerio Almeida, Anouk van Hooij, Nina Musch, Els Verhard, Suzanne van Veen, Louise Pierneef, Danielle de Jong, Raquel da Silva Corrêa, Thiago Thomaz Mafort, Rogério Rufino, Cristiana Santos de Macedo, Szymon M. Kiełbasa, Paul L.A.M. Corstjens, Luciana Silva Rodrigues, Annemieke Geluk

**Affiliations:** Dept. Infectious Diseases, Leiden University Medical Center, The Netherlands; Center for Technological Development in Health, Fiocruz, Rio de Janeiro, Brazil; Dept. Cell and Chemical Biology, Leiden University Medical Center, The Netherlands; Laboratory of Immunopathology, Faculty of Medical Sciences, Rio de Janeiro State University, Rio de Janeiro (UERJ), Rio de Janeiro, Brazil; Dept. of Pulmonary and Tisiology Care, Pedro Ernesto University Hospital (HUPE), Faculty of Medical Sciences, UERJ, Rio de Janeiro, Brazil; Dept. Biomedical Data Sciences, Leiden University Medical Center, The Netherlands

**Keywords:** pleural TB, Brazil, tuberculosis, diagnosis, biomarkers, UCP-LFA, cytokines, transcriptomics, RNA, microfluidic qPCR

## Abstract

Current diagnosis of pleural tuberculosis (PLTB) is based on highly invasive procedures. Therefore, blood-based host biomarkers could represent a low-invasive diagnostic alternative. Plasma and pleural effusion, as well as blood-and pleural fluid mononuclear cells (PBMCs and PFMCs), were sampled from patients with PLTB and other pleural diseases (OPLDs). In pleural effusion, ApoA1, C1q, CRP, IL-6, IFN-γ, IP-10, MIG, S100A12, SAA1/A2, and serpin-A3 were significantly higher in PLTB compared to OPLD, whereas only SAA1/A2 showed discriminatory potential in plasma. Increased mRNA levels were observed in PFMCs of PLTB for *CD8A*, *GBP5*, *SLAMF7*, *CXCL10*, *IL2*, *GNLY*, *IL23A*, *PDCD1* and *BCMA*, whereas *HMOX1*, *CD163*, *DUSP3*, *IGF1*, *GUSB* and *MARCO* were decreased. In PBMCs of PLTB, only *CCL22* expression was decreased. *GBP5* was significantly higher expressed in PLTB for both cell types.

This study shows the potential of transcriptomic and proteomic host biomarkers to differentiate PLTB from OPLDs also when applying low-invasive methods.

## Introduction

Tuberculosis (TB) remains one of the leading causes of death worldwide, caused by a single infectious pathogen, *Mycobacterium tuberculosis* (*Mtb*)^1^. In 2022, the World Health Organization (WHO) estimated that 10.6 million people were affected by TB. Moreover, an estimated 1.13 million people succumbed to TB among HIV-negative patients, and 167.000 deaths were reported among HIV-positive TB patients^1^. Currently, it is estimated that a quarter of the world’s population is latently infected with *Mtb*, and that 5 to 10% of these will develop active TB during their lifetime^1,2^. The risk of TB development increases with comorbidities such as malnutrition, smoking, diabetes, HIV infection, and alcoholism, as well as poor living conditions^1,3^. According to a cohort study on 100 million Brazilians, including 420,854 household contacts and 137,131 TB patients, the disease incidence among household contacts is 16 times higher than the general population^3^.

Pulmonary TB, typically characterized by involvement of the lungs, is the most frequently occurring and most contagious clinical form of TB^1^. However, about 20 to 25% of TB cases may manifest extrapulmonary involvement affecting other sites of the body. Amongst extrapulmonary forms of TB, pleural TB (PLTB) is the most common form in immunocompetent patients^4^.

Characteristic clinical symptoms of PLTB are fever, pleuritic chest pain due to fluid accumulation into the pleural cavity and nonproductive cough, impeding the use of sputum as a reliable clinical sample for diagnostics as in pulmonary TB^5^. Instead, PLTB diagnosis is based on the performance of a percutaneous or thoracoscopic pleural biopsy and the detection of *Mtb* in pleural effusion or pleural fragment samples^6^. However, due to its paucibacillary nature, sensitivity for detecting of *Mtb* in pleural effusion is disappointingly low^7^. A pleural biopsy which represents an even more invasive procedure, costly, and risky diagnostic approach requiring highly trained staff, could contribute to confirm TB diagnosis by the identification of the presence of caseating granuloma with or without acid-fast bacilli on histological specimens. Pleural effusions, characterized by accumulation of fluid between the parietal and visceral pleura, are, however, also present in malignant tumors, cardiac diseases, nontuberculous empyema, autoimmune diseases, and other pathologies^8^. Therefore, diagnostic tests based on low-invasive clinical samples that could differentiate PLTB from other pleural pathologies causing pleural effusion would contribute enormously to streamlining PLTB diagnostics.

Elevated levels of host proteins, such as inflammatory proteins IP-10, IFN-γ, and TNF^9,10^ have been observed in PLTB compared to other pleural diseases (OPLDs). These proteins have been evaluated as complementary biomarkers for the diagnosis of PLTB using, lateral flow assay based on up-converting phosphor reporter particles (UCP-LFA)^10^ or by flow cytometry (IFN-γ, TNF)^9^. Currently, host proteins such as adenosine deaminase (ADA), an enzyme secreted mainly by T lymphocytes and macrophages^4^, and IFN-γ by CD4^+^ and CD8^+^ T cells^11^, are mainly used as adjunct diagnostic tools for PLTB^4^. However, high levels of ADA are far from specific as they are also associated with other diseases, such as rheumatoid arthritis, lymphomas, malignant pleural effusions, and empyema^12,13^.

Host transcriptomics has been extensively shown to be an effective, exploratory approach to identify transcriptomic biomarkers for pulmonary TB^14^. This has led e.g. to the identification of RISK6 (*GBP2*, *FCGR1B*, *SERPING1*, *TUBGCP6*, *TRMT2A*, and *SDR39U1*), a transcriptomic biomarker signature for pulmonary TB that is also applicable for treatment response monitoring ^15,16^. Furthermore, the Cepheid GeneXpert MTB-HR comprising three host mRNA transcripts (3HR) test has been developed based on RNA expression of three genes (*KL2*, *DUSP3*, and *GBP5*). This 3HR test can differentiate patients with pulmonary TB from patients with other respiratory diseases (ORDs), with a sensitivity of 87% and specificity of 94%, regardless of HIV infection mainly on the African continent^17^. Also, in a tertiary care center in China^18^, the 3HR test discriminated TB from ORDs with a sensitivity of 82% and specificity of 66% ^18^.

Studies on biomarkers for PLTB using transcriptomics are scarce. Still, one study reported that, in a Brazilian cohort several genes could distinguish PLTB from non-TB patients^19^, as it showed that in PTLB patients, *ANKR22*, *GPB2* and *STAT1* mRNAs were higher compared to OPLD patients in pleural effusion samples as well as peripheral blood whereas *C1QB* was higher in the non-TB group^19^. Also, compared to lung diseases, such as pneumonia, and lung or metastatic cancer patients, PLTB showed higher mRNA expression in pleural effusion related to Th17 immune response genes (mainly *RORA*, *RORC*)^20^.

The current study aims to assess the discriminatory performance for PLTB *versus* OPLD by using proteomic and transcriptomic host biomarkers previously identified for active pulmonary TB in blood. To accomplish this, we assessed host protein levels in plasma and pleural effusion by UCP-LFA, as well as mRNA expression by microfluidic qPCR in peripheral blood mononuclear cells (PBMCs) and pleural fluid mononuclear cells (PFMCs).

## Results

### 3.1. Cohort characterization

Patients with complaints of chest pain due to pleural effusion and with other clinical indications for thoracentesis were recruited into this study (Table S1). These included 26 (39%) patients with TB pleural effusions, among which, three (12%) were HIV-positive. The remaining 40 (61%), with pleural effusions due to other pathologies, were included in the OPLD group. The average age for PLTB patients was 39, and 68 for OPLD patients. In the OPLD group, one individual presented with chylothorax, one with cirrhosis, one with lupus, two with empyema, four individuals with cardiovascular disease, five with unknown diagnosis, and 26 individuals with cancer. Among those classified with cancer, 8 individuals are classified with adenocarcinoma, 7 cancers with unknown primary site, 5 with pulmonary cancer, three with lymphoma, one ovary cancer, one with carcinoma and one breast cancer (Table S1). A significant difference in the age between PLTB and OPLD was observed, with the median of 39 and 67, respectively. Regarding the characteristics of the pleural fluid, a significant difference was observed related to ADA, the total of cells, total proteins, and lactate dehydrogenase (LDH) in pleural fluid between the PLTB and OPLD group. In contrast, glucose levels were significantly higher in OPLD patients.

### 3.2. Detection of host-derived proteins associated with PLTB diagnosis in pleural effusion and plasma

Since the user-friendly UCP-LFA platform was previously shown to be a practical rapid tool to detect disease-specific (inflammatory) biomarkers for leprosy, pulmonary TB, and COVID-19 ^10,21,22^, we used UCP-LFAs for targeted proteomic analysis of 15 host proteins, in pleural effusion and plasma of patients with PLTB and OPLDs^21–29^. Levels of ApoA1, C1q, CRP, MIG, IFN-γ, IL-6, IP-10, S100A12, SAA1/A2, and serpin-A3 in pleural fluid were significantly increased in PLTB compared to OPLDs whereas CFH, ferritin, IL-1Ra, NCAM, and TTR could not discriminate the two groups (Figure 1 and Figure S1). In contrast, when using plasma samples, only SAA1/A2 showed discriminatory potential between PLTB and OPLDs (Figure 2), whereas the other 14 showed similar levels in both patient groups (Figure S2). No significant differences were found regarding protein levels between males and females in pleural effusion and plasma samples (Figure S1 and S2).

**Figure 1:**
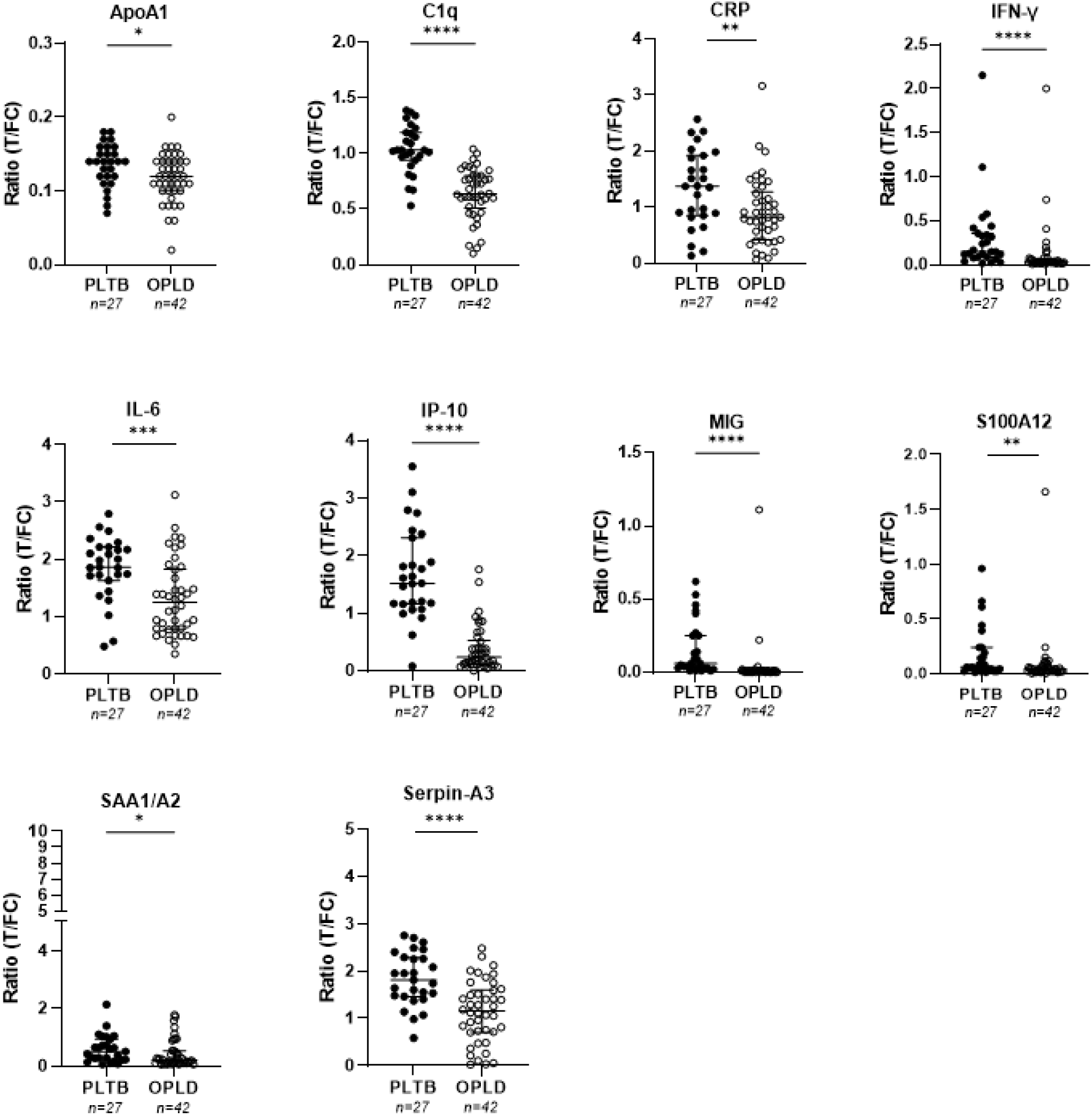
Detection of significantly different levels of host-derived proteins in pleural fluid of patients with PLTB (n=26) or OPLD (n=40). ApoA1, C1q, CRP, IFN-γ, IL-6, IP-10, MIG, S100A12, SAA1/A2 and serpin-A3 levels were measured by UCP-LFAs. Mann-Whitney U tests were performed to determine the statistical significance between groups (adjusted p-values: *p<0.05, **p<0.01, ***p<0.001, **** p<0.0001). The horizontal lines indicate the median with the interquartile range. T: test line; FC: flow control line. PLTB: Pleural Tuberculosis; OPLDs: other pleural diseases.

**Figure 2:**
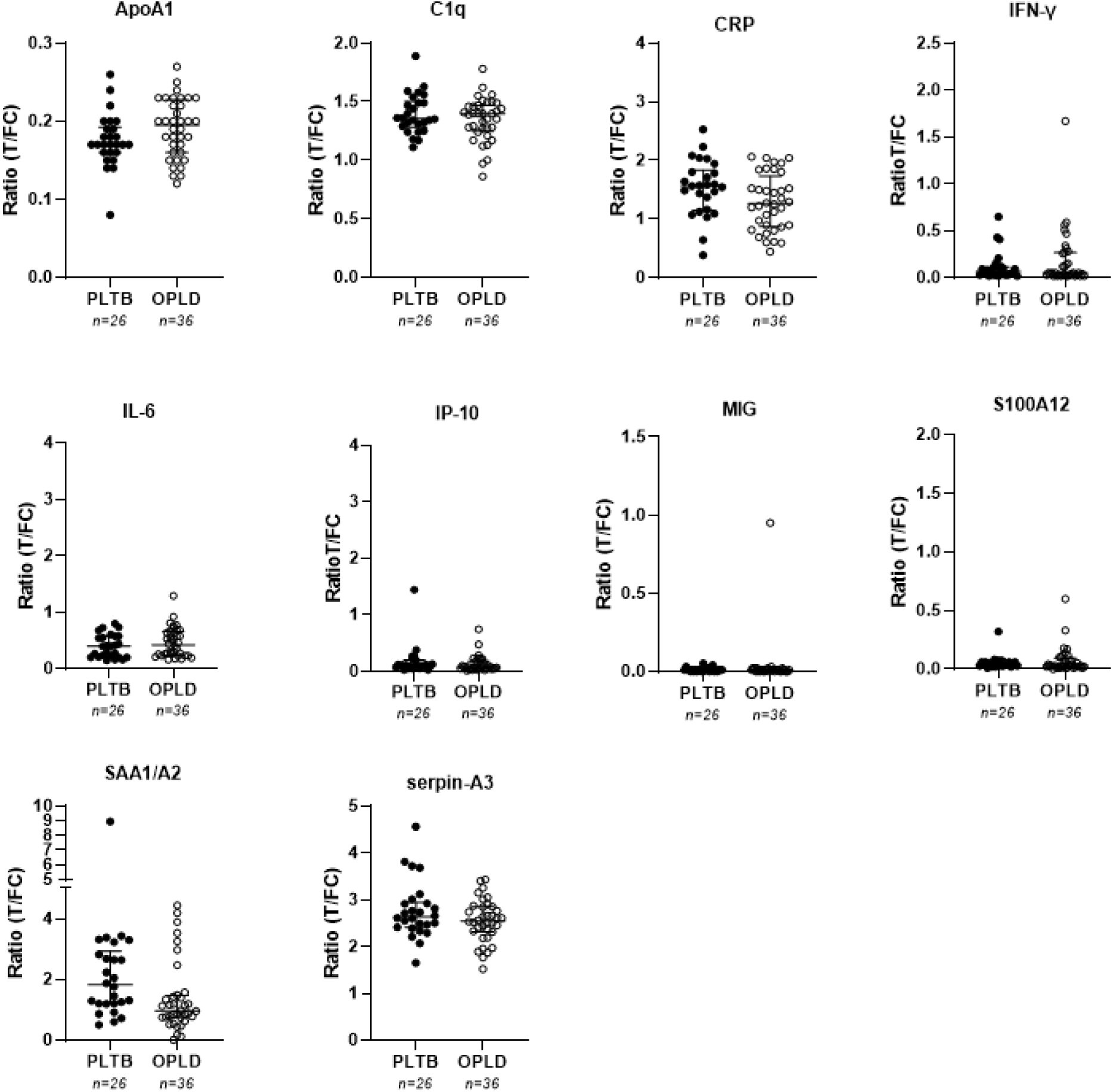
Host-derived protein levels in plasma of patients with PLTB (n=26) or OPLD (n=36). ApoA1, C1q, CRP, IFN-γ, IL-6, IP-10, MIG, S100A12, SAA1/A2 and serpin-A3 levels were measured by UCP-LFAs. Mann-Whitney U tests were performed to determine the statistical significance between groups (adjusted p-values: *p<0.05, **p<0.01, ***p<0.001, **** p<0.0001). Horizontal lines represent the median with the interquartile range. T: test line; FC: flow control line. PLTB: Pleural Tuberculosis; OPLDs: other pleural diseases.

### 3.3. Gene expression in PFMCs and PBMCs from microfluidic qPCR

To identify transcriptomic biomarkers for PLTB, high-throughput microfluidic qPCR was used to assess mRNA expression levels of 48 genes (Table S2) in PFMCs and PBMCs of patients with PLTB and OPLDs.

In PFMCs, it was observed that mRNA levels of *CD8A*, *GBP5*, *SLAMF7*, *CXCL10*, *IL-2*, *GNLY*, *IL-23A, PDCD1* and *BCMA* were significantly higher for PLTB than OPLDs (Figure 3). In contrast, *HMOX1*, *CD163*, *DUSP3*, *IGF1*, *GUSB,* and *MARCO* mRNA levels were significantly decreased in patients with PLTB compared to those with OPLDs (Figure 3). The other 32 genes measured in PFMCs were similarly expressed in both patient groups (Figure S3).

**Figure 3:**
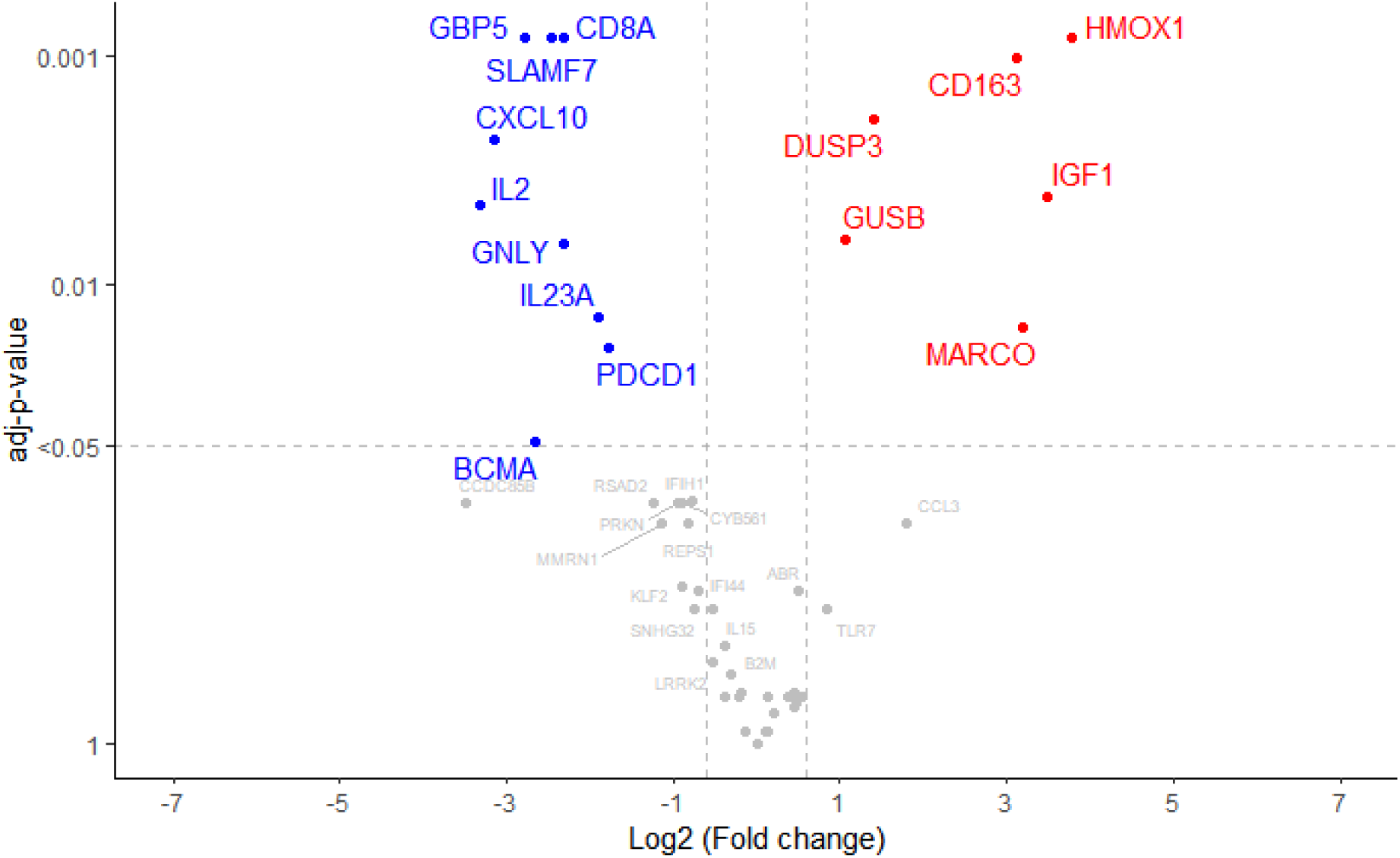
Volcano plot showing differential gene expression in PFMCs between PLTB and OPLDs. Logarithm of adjusted p-values is provided on the y-axis. Gene expression is provided on the x-axis as log_2_ fold change (FC) of OPLD compared to PLTB patients. Blue dots represent genes that are significantly (adjusted p-value <0.05 and log2FC < -1) higher expressed in PLTB. Red dots represent genes significantly (adjusted p-value <0.05 and log2FC > 1) lower expressed in PLTB. Gray dots represent genes that are not considered to be significantly different expressed between PLTB and OPLD.

For PBMCs, mRNA levels of *GBP5* were significantly higher in the PLTB group compared to OPLDs, whereas the reverse was found for *CCL22* (Figure 4). mRNA levels of none of the other genes varied significantly between these patient groups (Figure S4). No significant differences were found in mRNA levels between male and females in either PFMC or PBMCs (Figure S3 and S4).

**Figure 4:**
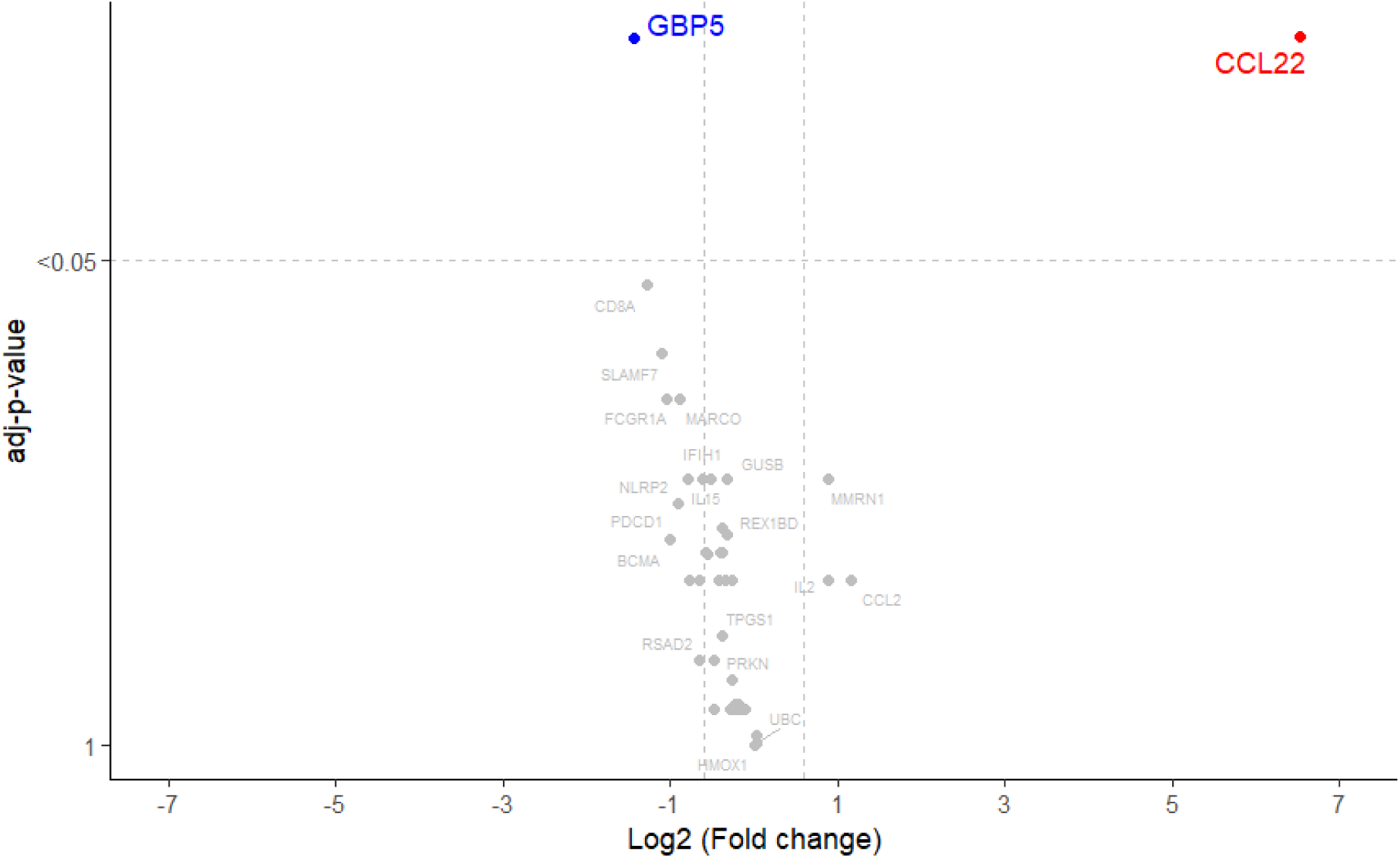
Volcano plot showing differential gene expression in PBMCs between PLTB and OPLDs. Logarithm of adjusted p-values is provided on the y-axis. Gene expression is provided on the x-axis as log_2_ fold change (FC) of OPLD compared to PLTB patients. Blue dots represent genes that are significantly (adjusted p-value <0.05 and log2FC < -1) higher expressed in PLTB. Red dots represent genes significantly (adjusted p-value <0.05 and log2FC > 1) lower expressed in PLTB. Gray dots represent genes that are not considered to be significantly different expressed between PLTB and OPLD.

Thus, for both PFMC and PBMC, mRNA levels of *GBP5* were significantly increased in the PLTB group compared to OPLDs (Figures 3 and 4). Furthermore, in PBMCs *CCL22* expression was significantly decreased, showing that these two genes represent promising transcriptomic biomarkers for PLTB using non-invasive samples.

To investigate whether the combination of blood-based transcriptomic and proteomic biomarkers could increase the diagnostic potential to discriminate PLTB from OPLD, cut-off levels for positivity, based on the optimal Youden’s index ^30^, for the genes *GBP5*, *CCL22* as well as the SAA1/A2 protein (Figure S5) were used to calculate the NUM-score. This score is the sum of the number of individual discriminatory biomarkers detected and is applied in diagnostic tests based on quantitative levels of multiple host biomarkers (Figure S6). This combined assessment of blood-based biomarkers resulted in improved sensitivity of 94,44% while sensitivity of the single biomarkers ranged from 61,11 % to 88,89 %. Specificity for the NUM-score was 82,14% which was lower than for GBP5 (89,29 %). However, GBP5 yielded a much lower concomitant sensitivity of 61,11% (Figure S6). As visualized in the Venn diagram (Figure S7), 7 out of 18 PLTB patients were positive for all 3 biomarkers, whereas 10 samples were either not detected with *GBP5*, *CCL22* mRNAs and SAA1/A2 protein. Based on this, the combination of blood-based biomarkers can provide improved diagnostic potential for PLTB than individual markers.

## Discussion

The current diagnosis of PLTB includes an initial assessment of the pleural cavity by ultrasound-guided thoracentesis, which is also considered a therapeutic approach to alleviate dyspnea, however, it is an extremely invasive and risky procedure. Due to the paucibacillary nature of PLTB, sensitivity for the detection of *Mtb* in pleural effusion or even in pleural fragments is low^7^. Therefore, a sensitive, non-invasive alternative method for detecting PLTB at point-of-care would be an innovation that could accelerate diagnosis and reduce patient distress.

In the current study, we focused on previously identified biomarkers for pulmonary TB using UCP-LFAs for 15 host proteins and microfluidic qPCR for 48 genes to identify, respectively, proteomic and transcriptomic biomarkers that differentiate PLTB from OPLD patients. Compared to OPLD, the highly inflammatory nature of PLTB was predominantly reflected in pleural effusion and not in plasma. This confirmed previous findings for the presence of IP-10 in pleural effusion samples of PLTB patients in a Gambian^10^ and IFN-γ in a Brazilian population^9^. To identify biomarkers for PLTB that can be measured in low-invasive samples, we applied UCP-LFAs specific for host biomarkers previously identified for pulmonary TB^21,31–33^.

Our data is aligned with the characteristic “immune response compartmentalization” occurring during the development of PLTB and it is represented by the accumulation of Th1 pro-inflammatory molecules in pleural effusion rather than in peripheral blood^9^. The chemokines MIG and IP-10, of which high levels are present in pleural effusions of PLTB patients in our study, attract activated IFN-γ producing Th1 lymphocytes to the site of infection^9,34,35^. Previous reports demonstrated that both MIG and IP-10 were present at higher levels in PLTB compared to OPLDs patients^35,36^. Also, in this study, a series of inflammatory mediators, such as CRP, IL-6, C1q, serpin-A3, and S100A12, were detectable^9,37–39^. S100A12, which has been described to have antimicrobial activity for both *Mtb* and *M. leprae*^40^, and serpin-A3, a regulatory enzyme involved in physiological functions like wound healing and apoptosis^41^ protecting from tissue damage during active TB^41^, were present in pleural effusions in PLTB patients. Since ApoA1 can reduce the production of pro-inflammatory molecules once PLTB progresses^42,43^, the higher concentration of ApoA1 detected in PLTB could be due to its anti-inflammatory function once PLTB progresses to dampen a highly inflammatory phase in the pleural cavity^9,38^.

The host protein SAA1/A2 was significantly increased in pleural fluid samples and also demonstrated discriminatory potential in plasma in PLTB compared to OPLD patients. This protein is a member of the human serum amyloid A protein (SAA) family, reflecting the inflammatory state of the lung in PLTB. Thus, this shows the potential of SAA1/2 as a biomarker for PLTB mainly in pleural fluid, with a trend for higher levels in PLTB in blood samples that have an essentially lower invasive character. These findings are in line with increased SAA1/A2 levels detected in plasma of pulmonary TB patients^21,31,44^. Applying fingerstick blood^45^ for detection of SAA1/2 using the low-complexity UCP-LFA platform could provide a rapid (adjunct) diagnostic tool for PLTB that can be performed at point-of-care thereby avoiding the need for percutaneous or thoracoscopic pleural biopsy^45–47^.

Our search for transcriptomic biomarkers for PLTB led to the detection of higher mRNA levels of *BCMA*, *CD8A*, *GNYL*, *GBP5*, *IL2*, *IL23A*, *CXCL10*, *PDCD1* and *SLAMF7* in PLTB than in OPLDs. *SLAMF7* has an essential role in the activation of cells such as CD8 T lymphocytes^48^. It is also related to T lymphocyte exhaustion, where its activation leads to the expression of inhibitory receptors such as PD-1 and CTLA-4^49^. The increased expression of *IL2*, *IL23A* and *CXCL10* confirm the pro-inflammatory nature of the response against *Mtb*, which is in line with the proteomics data of this present study. On the other hand, mRNA levels of *CD163*, *DUSP3*, *GUSB*, *HMOX1*, *IGF1*, and *MARCO* were decreased in PFMCs of the PLTB group compared to the OPLD group. CD163 is a specific biomarker for M2 (anti-inflammatory) macrophages^50^. In contrast to our results, *CD163* was found to be higher in tissue samples of patients with pulmonary tuberculoma with high inflammatory status when compared to those with moderate inflammatory status^51^. This could likely be because tissue may harbor macrophages with anti-inflammatory profile, whereas PFMCs are characterized by a highly inflammatory environment with the absence of this cell with an anti-inflammatory profile.

*GBP5*, which plays a vital role in the assembly of AIM2 and NLRP3 inflammasomes^52,53^, has been extensively described as a transcriptomic biomarker to differentiate TB patients from LTBI, other pulmonary diseases, and healthy individuals^54–56^. Our findings have shown that expression of *GBP5* was increased in PLTB in both PBMCs and PFMCs, indicating the potential of this gene as a biomarker for differentiating PLTB from other pathologies, even using low-invasive blood samples.

Recently, the use of *GBP5* expression as a biomarker to distinguish active TB from ORDs using fingerstick blood test was described^17^ as part of interim results of Cepheid GeneXpert MTB-HR prototype using capillary blood analysis of 195 participants from four countries. Besides *GBP5*, which was increased in TB compared to ORDs patients, this test included two other genes: *DUSP3*, which was also higher in TB patients and *KLF2* which was similar in TB and OPLD^17^. Based on the expression of these genes, a TB score ((Ct GBP5 + Ct DUSP3)/2 – Ct KLF2) was calculated^17^, providing discrimination between TB and OPLDs based on their differential gene expression in active TB patients. However, in our study, *DUSP3* expression was lower in PLTB than in OPLD prompting more evaluation in larger PLTB cohorts. Still, based on *GBP5* levels in PLTB, the Cepheid test would represent an excellent rapid test to diagnose PLTB in a clinical setting.

Finally, it was observed that *CCL22* mRNA levels were decreased in PBMCs of the PLTB group. This chemokine has an important chemotactic activity leading to recruitment of regulatory T lymphocytes^57,58^. This gene expression decrease could be due to the lack of regulatory response during the PLTB infection, characterizing the high inflammatory status. It was described that IL-26 induces the increase of CCL22 levels in mononuclear cells isolated from pleural effusions of PLTB patients^59^. In contrast to our results, it was observed that individuals with active TB HIV+ displayed a transcriptomic signature with increased mRNA levels of *CCL22* and other genes such as *FCGR1A, TIMP2* and *TNFRSF1A*, and decreased *IL-7R* and *CD8A* expression compared to patients with non-active TB HIV+^60^.

In summary, proteomic and transcriptomic biomarkers were identified using UCP-LFA and qPCR, respectively. Both biomarker profiles confirmed the inflammatory character of the pleural cavity in PLTB. Moreover, only SAA1/A2 protein levels showed a discriminatory potential in pleural effusion and plasma of PLTB patients compared to those with OPLDs, indicating that SAA1/A2 levels in plasma reflect the inflammatory state of the pleura.

The mRNA levels of *GBP5* were higher in both PBMCs and PFMCs of PLTB patients, while only *CCL22* was significantly lower expressed in PBMCs of PLTB. Therefore, these genes, like the host protein SAA1/A2, represent promising biomarkers to distinguish PLTB from OPLD, even in low-invasive blood samples.

## Limitations of the study

Due to strict inclusion criteria, a relatively low number of PLTB patients was recruited into this study. Also, to ensure global applicability, similar studies should be performed in other endemic areas, besides this Brazilian cohort. Furthermore, it should be noted that the cohort studied concerned outpatients before the COVID-19 pandemic, and therefore, SARS-CoV-2-infected patients were not included in the OPLD group. Thus, we cannot exclude that (future) severe viral diseases leading to pleural effusions at diagnosis or with clinical indications for thoracentesis, provide a similar biomarker pattern. Finally, to allow quantitative transcriptomic biomarker comparison between blood and pleura, PBMCs and PFMCs were respectively used for mRNA expression analysis, which excludes gene expression in other blood cells that may contribute to the production of other valuable biomarkers.

## Author contributions

Conceptualization: AG

Data curation: MA, SV, EV, NM, AH, DJ, LP, RC, LR

Formal analysis: MA, SV, SK

Funding acquisition: PC, CM, LR, AG

Investigation: MA, EV, SV, AH, DJ, LP, RC, TM, RR

Methodology: MA, SV, PC, AG

Project administration: AG

Resources: PC, LR, AG

Software: MA, SV, SK

Supervision: CM, SK, AG

Validation: MA, SK

Visualization: MA, SK, AG

Writing – original draft: MA, LR, AG

Writing – review and editing: MA, EV, SV, CM, NM, LP, RC, LR, AH, PC, SK, AG

## Acknowledgements

This work was made possible thanks to the Q.M. Gastmann-Wichers foundation (AG) and LUMC Global (MA). This study was supported by Fundação Oswaldo Cruz (FIOCRUZ) and Fundação para o Desenvolvimento Científico e Tecnológico em Saúde (FIOTEC). The authors wish to thank all patients and staff from the Pedro Ernesto University Hospital, in Rio de Janeiro, Brazil. The graphical abstract for this article was created with <u>BioRender.com</u>.

## Subject details

### Ethics statement

Ethical approvals for this observational and cross-sectional cohort study were received from the institutional ethics approving committee at Pedro Ernesto University Hospital (HUPE), Rio de Janeiro State University (UERJ), Rio de Janeiro, RJ, Brazil (approval number 1.100.772) at 10/06/2015. Participants were informed about the study objectives. All participants agreed to participate in the study and signed a written informed consent before inclusion. All participants were informed about their right to refuse to take part or withdraw from the study without consequences. All samples were fully anonymized before processing to protect the study participants’ identities.

### Patient recruitment

Between June 2016 and November 2019, patients were enrolled at the Pedro Ernesto University Hospital, a referral center for various tertiary care unit at the Rio de Janeiro State University, Brazil. Patients who presented clinically and with X-rays suggestive of undiagnosed pleural effusions at diagnosis or with clinical indications for thoracentesis were eligible for recruitment into the study. Patients who had been taking anti-TB drugs for more than seven days, who were pregnant, under 18 years of age, or who refused consent were not included in the study. An HIV test was offered to all patients included in the study. Blood and pleural effusion samples were collected before treatment.

A total of 66 patients, including 34 men and 32 women aged between 18 and 92 years old (table S1), were included. Among these, 26 were diagnosed with TB pleural effusions and 40 with pleural effusions due to other pathologies such as chylothorax, lupus, cardiovascular disease, empyema, cirrhosis, cancer, and pleural effusion undiagnosed.

### Diagnosis criteria

Radiological findings were based on chest X-rays and classified as unilateral or bilateral pleural effusions. The diagnosis of pleural TB was confirmed when at least one of the following criteria was met: 1) a positive result for acid-fast bacillus (AFB) staining, or Xpert MTB/RIF of pleural effusion; 2) a positive result for Lowenstein-Jensen (LJ) medium culture of pleural effusion or pleural tissue; or 3) detection of caseous granuloma by histopathological analysis in pleural tissue ^61^. Patients presenting with signs and symptoms, such as fever, dyspnea, cough, chest pain and night sweats for > 3 weeks who had adenosine deaminase (ADA) > 40IU/L in pleural fluid, negative microbiological and histopathologic examinations, but improved after anti-TB treatment, were classified as pleural TB cases^62^.

The patients were analyzed for ADA, lactic dehydrogenase, glucose, cholesterol, amylase, protein, and albumin concentrations. Pleural effusion was additionally tested for total and differential cell count, and Gram staining. The pleural effusion was considered exudative when the pleural effusion/serum protein ratio (R) was higher than 0.5 and mononuclear when it contained more than 50% of mononuclear cells^63^. Pleural effusion aliquots were collected by a trained pulmonologist by ultrasound-guided thoracentesis which were separated, numbered, and frozen at -80°C. Subsequently, four fragments of parietal pleura were obtained with a Cope needle: three for histopathologic examination and one for LJ culture.

## Method details

### Sample preparation and isolation of cells

Ultrasound-guided thoracentesis was performed for the collection of peripheral blood mononuclear cells (PBMCs) in heparin tubes, which were obtained at > 92% purity from venous blood using a density gradient (Ficoll-Hypaque, Sigma-Aldrich Inc., USA) as described^64^. Pleural fluid mononuclear cells (PFMCs) were obtained by directly centrifuging pleural effusions at 500 g for 8 min at 25°C^11^. PFMCs were washed three times with endotoxin-free phosphate-buffered saline (PBS 50 mM, pH 7.2, Sigma-Aldrich) at 500 g for 8 min at room temperature (RT), and 4°C in the last wash. To the PBMCs and PFMCs (5 x 10^6^ cells each), 1 mLof TriZol (Life Technologies, Grand Island, NY, USA) was added, and samples were frozen at -80°C until RNA isolation. Plasma and pleural effusion of the samples related to blood and pleural effusion, respectively, were collected and frozen at -80 °C.

### RNA isolation

Total RNA was extracted from PBMCs and PFMCs using TriZol reagent (Invitrogen, Thermo Fisher Scientific, Waltham, USA). RNA isolation was performed according to the manufacturer’s protocol. RNA concentrations were measured by spectrophotometry using a DS-11 DeNovix spectrophotometer (DeNovix Inc., Wilmington, Delaware, USA)^65^. For RNA expression analyses, samples were diluted to 50 ng/µL.

### UCP Conjugates

The UCP-LFA was developed and carried out to enable the quantitative detection of the desired biomarker^22,24,25,29,66,67^. UCP nanomaterials were provided by Intelligent Material Solutions Inc. (Princeton, New Jersey, USA). The UCP-LFA strips for interleukin-6 (IL-6), interleukin-1 receptor antagonist (IL-1Ra), complement component 1q (C1q), C-reactive protein (CRP), interferon gamma-induced protein 10 (IP-10), S100 calcium-binding protein A12 (S100A12), serum amyloid A1 and A2 (SAA1/A2), apoA1, and ferritin were produced as described previously^21–23,28^. The UCP conjugates for chemokine monokine induced by gamma interferon (MIG) and complement factor H (CFH) were prepared with mouse anti-human MIG (49106, R&D systems, Minneapolis, USA), and mouse anti-human CFH (MAB4779, R&D systems), respectively, at a concentration of 50 µg antibody per mg UCP. The UCP conjugates for neural cell adhesion molecule (NCAM/CD56) and serpin-A3 were prepared with goat anti-human NCAM Ab (AF2408, R&D systems) and goat antihuman serpin-A3 (AF1295, R&D systems), respectively, at a concentration of 50 µg antibody per mg UCP. UCP strips for transthyretin (TTR) were prepared with rabbit anti-human TTR (EPR20073-155, ABCAM) at a concentration of 50 µg antibody per mg UCP. The UCP conjugates for interferon-γ (IFN-γ) were prepared with mouse anti-human IFN-γ (B-B1, Medix Biochemica) at a concentration of 25 µg antibody per mg UCP.

### Lateral flow Strips

Lateral flow (LF) strips were produced as described previously^21–29^. Details concerning strips for additional UCP-LF assays not previously described (CFH, IFN-γ, MIG, NCAM, serpin-A3, and TTR) are summarized below. For UCP-LFA strips for MIG and CFH have a test (T) line comprised of a mouse-anti-human MIG and mouse-anti-human CFH (AF392, AF4779, R&D systems), respectively. For NCAM and serpin-A3 the T lines comprise goat-anti-human NCAM and goat-anti-human serpin-A3 (AF2408, Clone 213907, R&D systems), respectively. The T line for TTR comprised rabbit-anti-human TTR (AB77905; ABCAM), and for IFN-γ, mouse anti-human IFN-γ (B-G1, Medix Biochemica). All T lines contained 200 ng antibody per 4 mm of width. The flow control (FC) lines for the MIG, IFN-γ and CFH comprised a goat-anti-mouse IgG antibody (M8642, R&D systems), for NCAM and serpin-A3, a rabbit-anti-goat IgG antibody (RαG; G4018, R&D systems) was used, and for TTR a goat-anti-rabbit IgG antibody (R4880, ABCAM). The TTR UCP reporter conjugate was applied to the sample/conjugate-release pad at a density of 200 ng per 4 mm width in a buffer containing 5% (w/v) sucrose, 50 mM Tris pH 8.0, 0.6 mM Borate pH 8, 135 mM NaCl, 0.5% (w/v) BSA, and 0.25% Tween-20. The pads were dried for 1 hour at 37 °C. Until used, LF strips were stored at ambient temperature in plastic containers with silica dry pads.

### Detection of host proteins by UCP-LFAs

Both plasma and pleural effusion samples were diluted in high salt finger stick (HSFS) buffer (100 mM Tris pH 8, 270 mM NaCl, 1% (w/v) BSA, and 1% (v/v) Triton X-100) as follows: 1:10 (for MIG, IFN-γ, IL-1Ra, NCAM, IP-10, IL-6, and ferritin), 1:100 (for S100A12), 1:1000 (for CFH, serpin-A3, CRP, SAA1/A2, and TTR), 1:10000 (for C1q and ApoA1). The diluted plasma/pleural effusion samples (100 µL) were applied directly to the target-specific LF strips (IL-6, ferritin, S100A12, ApoA1, IP-10, CRP, SAA1/A2, and TTR), (dry format; UCP particles incorporated into the test strip^21–29^). For IL-1Ra, MIG, NCAM, CFH, serpin-A3, IFN-γ and C1q, 200 ng of target-specific UCP reporter was included with the 100 µL diluted plasma/pleural effusion before application to the target-specific LF strips (wet format; UCP particles were added into a plate well with the sample before the strips were inserted to the well). The LF strips were scanned to detect the UCP reporter (980 nm excitation and 550 nm emission, UPCON; Labrox, Finland). Results are presented as the ratio value between the T (test) over FC (flow control) signals based on relative fluorescence units (RFUs) measured at the respective lines (peak areas).

### RNA expression analysis

High-throughput microfluidic qPCR was performed using the Biomark HD system (Standard BioTools, South San Francisco, CA)^65^. Reverse Transcription Master Mix (Standard BioTools) was used to convert 50 ng of RNA into cDNA according to manufacturer’s instructions, as described previously^65^. Prior to real-time amplification with the 48.48 Dynamic Array™ integrated fluidic circuit (IFC), the cDNA was preamplified within 14 cycles to increase the concentration of material required for the RT-qPCR technique using Preamp Master Mix and a pool of limited amounts of all TaqMan assays according to the manufacturer’s instructions. Thermal cycling conditions were: 95°C for 2 minutes, followed by 14 cycles at 95°C for 15 seconds, and 60°C for 4 minutes. Preamplified cDNA was diluted 1:5 in a dilution reagent (10 mM Tris-HCL, pH 8.0, 0.1 mM EDTA) and stored at -20°C until further processing. The mRNA transcripts of 48 genes (Table S1) were measured by microfluidic qPCR using 48.48 IFC chips on the Biomark HD system (Standard BioTools), according to the manufacturer’s instructions. qPCR was performed with the Biomark HD using the following thermal cycling protocol: 95°C for 10 minutes, followed by 40 cycles at 95°C for 15 seconds and 60°C for 1 minute. Data was analyzed using Fluidigm Real-Time PCR Analysis Software (v. 4.5.2, Standard BioTools Standard BioTools). A cycle threshold (Ct) value ≤ 30 was determined as the cutoff for reliable detection. Relative target gene expression was determined by calculating ΔCt using GAPDH as a reference gene. Since mRNA levels of immune genes were expressed as ΔCt values, larger values represent lower mRNA levels in the sample. To obtain the correct direction, the ΔCt values were multiplied by -1.

### Statistical analysis

The comparison between the PLTB and OPLD regarding to the sociodemographic and clinical characteristics of the patients with pleural effusion was made using Fisher’s exact test for comparison of the relative frequencies of nominal/categorical variables, and nonparametric Mann-Whitney test for continuous variables. The difference in median expressions between groups of the UCP-LFA and the microfluidic qPCR was evaluated with the Mann-Whitney U test. Differences in protein levels between groups were considered significant when their corresponding p-values, adjusted for multiple testing with the Bonferroni-Holm method, were below the 0.05 threshold. The gene expression was considered significant when their corresponding p-values, adjusted for multiple testing with the Bonferroni-Holm method, were below the 0.05 threshold and above 1 or below -1 log2FC threshold. Results were expressed as a median ± IQR for each group for proteins analyses and genes expression is provided on the axis as log_2_ fold change (FC) of OPLD compared to PLTB patients. The analyses were performed using R Studio (version 4.0.2) and GraphPad Prism (version 9.3.1) for Windows.

