## Supplemental information for "Proteomic and transcriptomic host biomarkers for detection of pleural tuberculosis"

**Supplementary tables and figures**

**Supplementary Table 1: Characteristics of the study population**

| | **PLTB (n=26)** | **OPLDs (n=40)** | **p-value** |
|---|---|---|---|
| **Gender, n (M:F)** | 16:10 | 18:22 | 0.2163 |
| **Age (median in years)[a]** | 39 | 68 | <0.0001 |
| **HIV status** | | | |
| -positive | 3 | - | 0.0800 |
| -negative | 23 | 33 | |
| -unavailable | - | 7 | |
| **Pleural effusion characteristics** | | | |
| -exudate | 22 | 30 | 0.6363 |
| -transudate | 1 | 3 | |
| -unavailable | 3 | 7 | |
| **Pleural fluid – laboratory analysis[a]** | | | |
| - ADA, IU/L | 50.60 | 11.70 | <0.0001 |
| - total cells/mm$^3$ | 3600 | 1375 | 0.0026 |
| - mononuclear cells, % | 95 | 84.60 | 0.0739 |
| - polymorphonuclear cells, % | 5.0 | 15.45 | 0.0739 |
| - glucose, mg/dL | 78.00 | 105.0 | <0.0001 |
| - total proteins, g/dL | 5.2 | 4.6 | 0.0086 |
| - albumin, g/dL | 2.9 | 2.7 | 0.2792 |
| - LDH, IU/L | 306 | 171 | 0.0103 |
| **PLTB diagnostic criteria** | | | |
| - microbiology | 1 | - | |
| - histopathology | 10 | - | |
| - ADA > 40IU/L, clinical findings and full recovery after anti-TB treatment | 15 | - | |
| **Pleural effusion cause** | | | |
| -TB | 26 | - | |
| -Cancer | - | 26 | |
| -Cardiac | - | 4 | |
| -Chylothorax | - | 1 | |
| -Cirrhosis | - | 1 | |
| -Nontuberculous empyema | - | 2 | |
| -Lupus | - | 1 | |
| -undefined diagnosis | - | 5 | |

PLTB: Pleural tuberculosis.

OPLD: Other pleural diseases.

ADA: Adenosine deaminase.

LDH: Lactate dehydrogenase.

**Supplementary Table 2: Primers and probe Assay IDs.**

| Gene name | Ensembl ID | Thermofisher Assay ID |
|---|---|---|
| ***DUSP3*** | **ENSG00000108861** | Hs01115776_m1 |
| ***TPGS1*** | **ENSG00000141933** | Hs00293366_m1 |
| ***CYB561*** | **ENSG00000008283** | Hs00157220_m1 |
| ***MT-CYB*** | **ENSG00000198727** | Hs02596867_s1 |
| ***TLR7*** | **ENSG00000196664** | Hs00152971_m1 |
| ***IL-2*** | **ENSG00000100385** | Hs00174114_m1 |
| ***GBP5*** | **ENSG00000154451** | Hs00369472_m1 |
| ***UBC*** | **ENSG00000150991** | Hs00824723_m1 |
| ***GNLY*** | **ENSG00000115523** | Hs01120727_m1 |
| ***MT-ND2*** | **ENSG00000198763** | Hs02596874_g1 |
| ***CCL3*** | **ENSG00000277632** | Hs00234142_m1 |
| ***CXCL10*** | **ENSG00000169245** | Hs00171042_m1 |
| ***KLF2*** | **ENSG00000127528** | Hs00360439_g1 |
| ***GAPDH*** | **ENSG00000111640** | Hs99999905_m1 |
| ***HMOX*** | **ENSG00000103415** | Hs01110250_m1 |
| ***MT-ND4*** | **ENSG00000198886** | Hs02596876_g1 |
| ***CD163*** | **ENSG00000177575** | Hs00174705_m1 |
| ***IL-15*** | **ENSG00000134470** | Hs01003716_m1 |
| ***PDCD1*** | **ENSG00000188389** | Hs01550088_m1 |
| ***ABR*** | **ENSG00000159842** | Hs01077828_m1 |
| ***LRRK2*** | **ENSG00000188906** | Hs01115057_m1 |
| ***MT-ND5*** | **ENSG00000198786** | Hs02596878_g1 |
| ***IL-23A*** | **ENSG00000110944** | Hs00372324_m1 |
| ***FCGR1A*** | **ENSG00000150337** | Hs00174081_m1 |
| ***SLAMF7*** | **ENSG00000026751** | Hs00904275_m1 |
| ***GUSB*** | **ENSG00000169919** | Hs00939627_m1 |
| ***BCMA*** | **ENSG00000048462** | Hs00171292_m1 |
| ***REPS1*** | **ENSG00000135597** | Hs01016191_m1 |
| ***NLRP2*** | **ENSG00000022556** | Hs01546932_m1 |
| ***IFI44*** | **ENSG00000137965** | Hs00197427_m1 |
| ***CCDC85B*** | **ENSG00000175602** | Hs00255227_s1 |
| ***B2-M*** | **ENSG00000166710** | Hs00187842_m1 |
| ***IGF1*** | **ENSG00000017427** | Hs01547656_m1 |
| ***REX1BD*** | **ENSG00000006015** | Hs00215835_m1 |
| ***RSAD2*** | **ENSG00000134321** | Hs00369813_m1 |
| ***IFIH1*** | **ENSG00000115267** | Hs00223420_m1 |
| ***MMRN1*** | **ENSG00000138722** | Hs01113299_m1 |
| ***CCL22*** | **ENSG00000102962** | Hs01574247_m1 |
| ***CD8A*** | **ENSG00000153563** | Hs00233520_m1 |
| ***SNHG32*** | **ENSG00000204387** | Hs00382553_m1 |
| ***FABP9*** | **ENSG00000205186** | Hs00908573_m1 |
| ***OAS1*** | **ENSG00000089127** | Hs00973635_m1 |
| ***MT-CO1*** | **ENSG00000198804** | Hs02596864_g1 |
| ***PRKN*** | **ENSG00000185345** | Hs01038322_m1 |
| ***CCL2*** | **ENSG00000108691** | Hs00234140_m1 |
| ***TAOK3*** | **ENSG00000135090** | Hs00937694_m1 |

| | | |
|---|---|---|
| ***GZMB*** | **ENSG00000100453** | Hs00188051_m1 |
| ***MARCO*** | **ENSG00000019169** | Hs00198937_m1 |

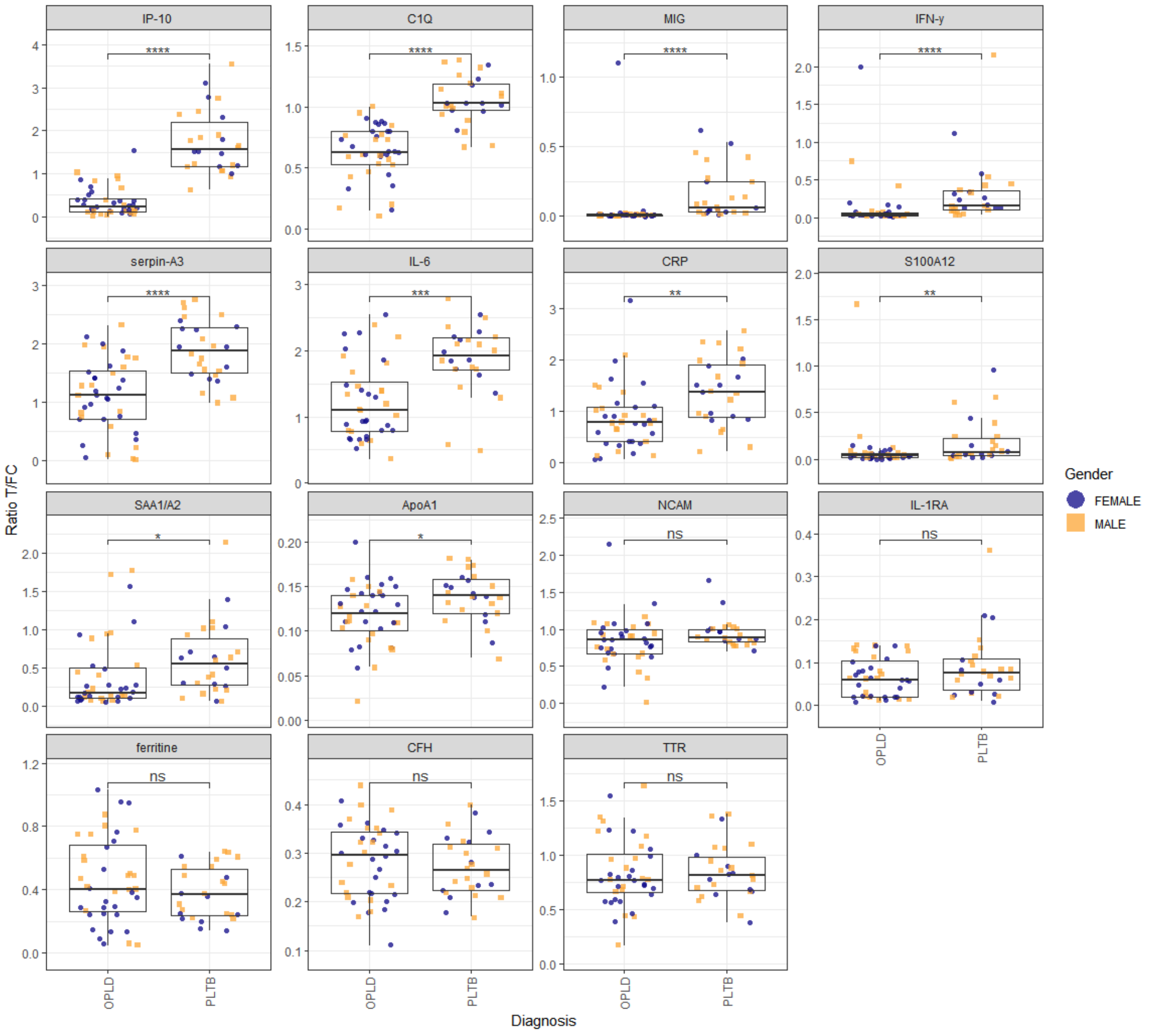


**Supplementary figure 1:** Detection of proteins in pleural fluid of patients with PLTB (n=26) or OPLDs (n=40). ApoA1, C1q, CFH, CRP, ferritine, IFN-y, IL-6, IL-1Ra, IP-10, MIG, NCAM, S100A12, SAA1/A2, serpin A3, and TTR levels were measured by UCP-LFAs. Adjusted p-values: *p<0.05, **p<0.01, ***p<0.001, **** p < 0.0001. T: test line; Fc: flow control line. PLTB: Pleural Tuberculosis; OPLDs: other pleural diseases.

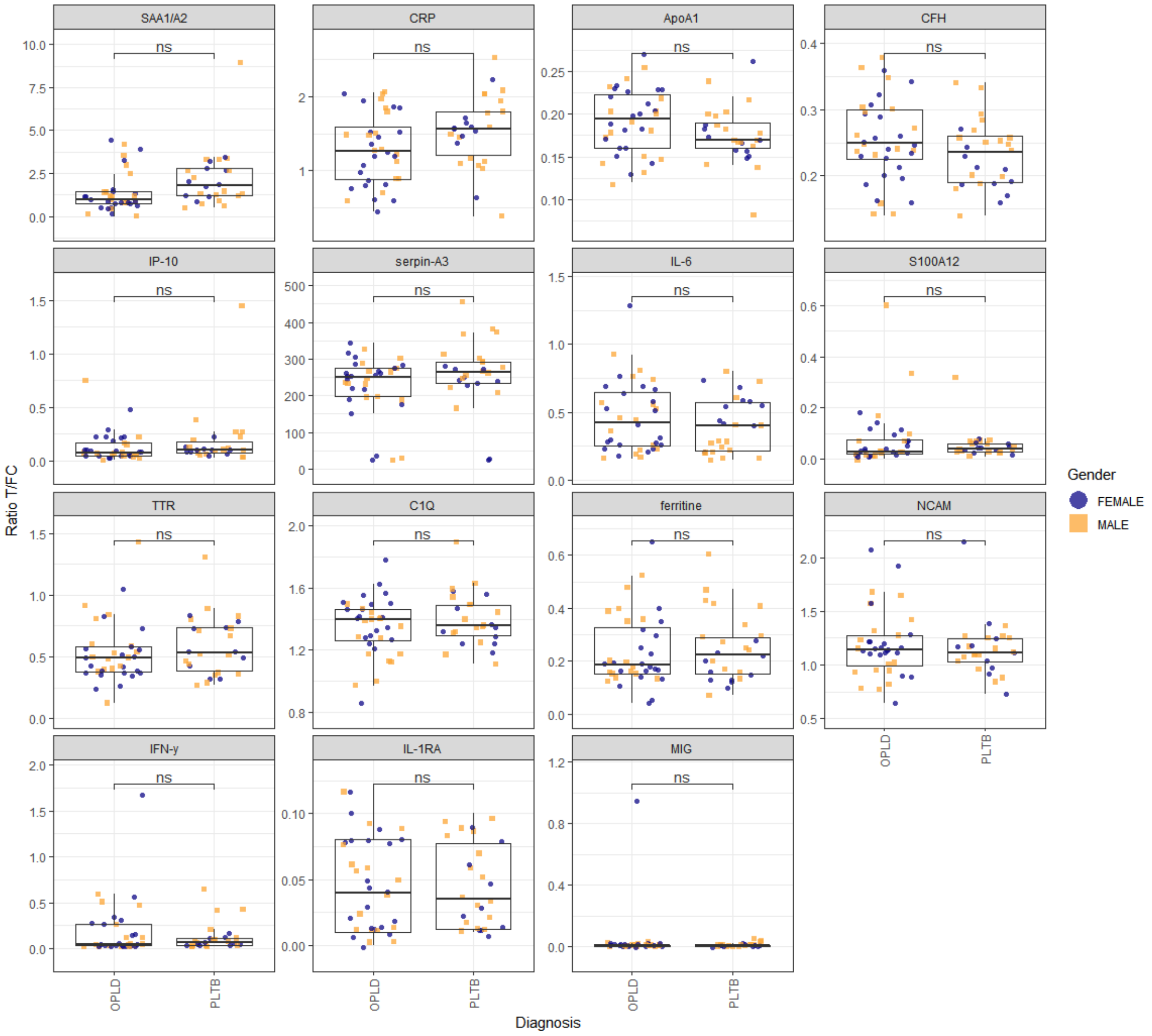


**Supplementary figure 2:** Detection of proteins in plasma of patients with PLTB (n=26) or OPLDs (n=36). ApoA1, C1q, CFH, CRP, ferritine, IFN-y, IL-6, IL-1Ra, IP-10, MIG, NCAM, S100A12, SAA1/A2, serpin A3, and TTR levels were measured by UCP-LFAs. Adjusted p-values: *p<0.05, **p<0.01, ***p<0.001, **** p < 0.0001. T: test line; Fc: flow control line. PLTB: Pleural Tuberculosis; OPLDs: other pleural diseases.

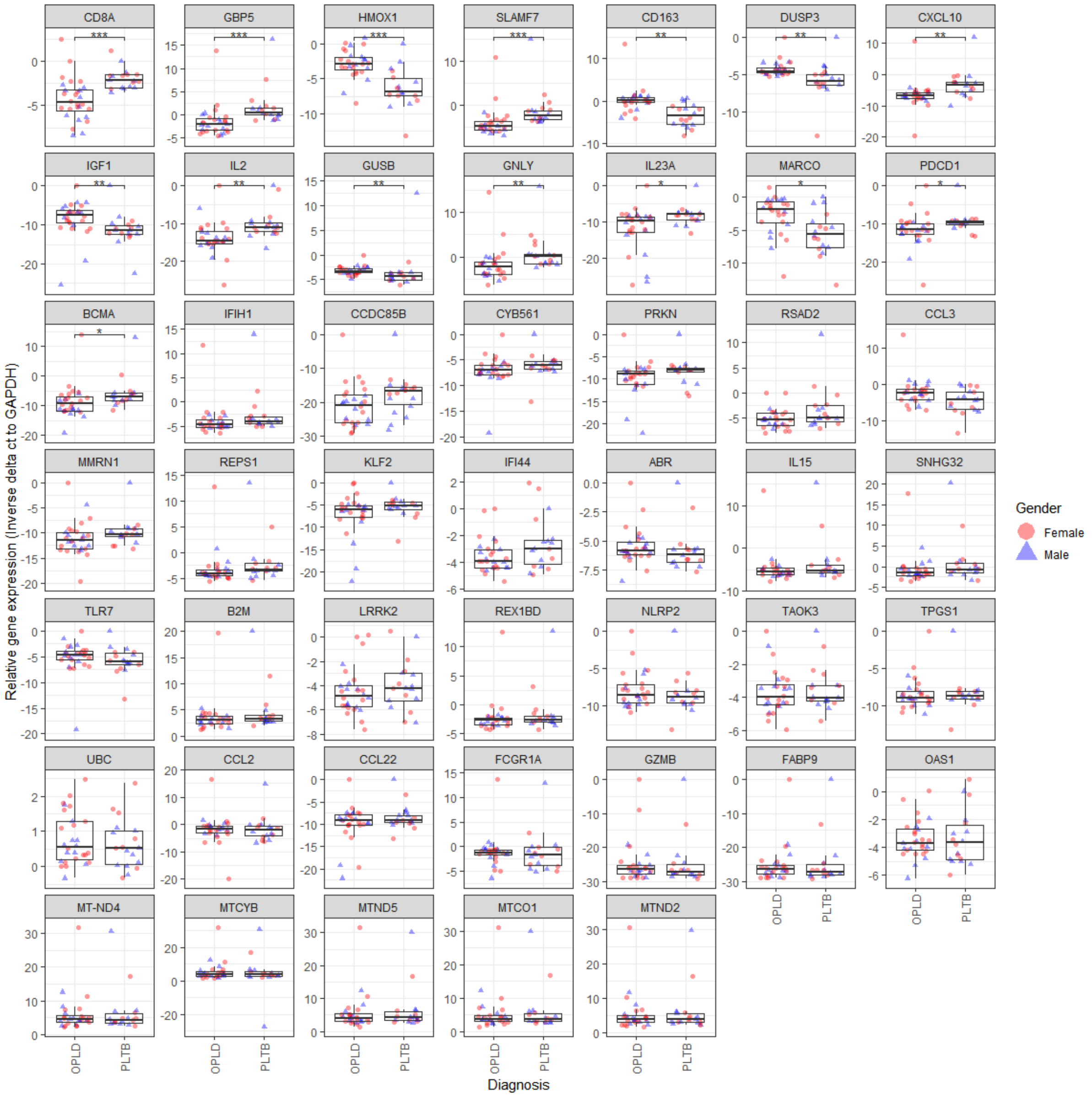

**Supplementary figure 3:** Gene expression in PFMC of patients with PLTB and OPLD. The mRNA levels were measured by microfluidic RNA-seq in human samples of patients with PLTB (n=18), OPLDs (n=28). Relative target gene expression was determined by calculating ΔCt using GAPDH as a reference gene. Since mRNA levels of immune genes were expressed as ΔCt values, larger values represent lower mRNA levels in the sample. For the purpose of obtaining the correct direction, the ΔCt values were multiplied by -1.

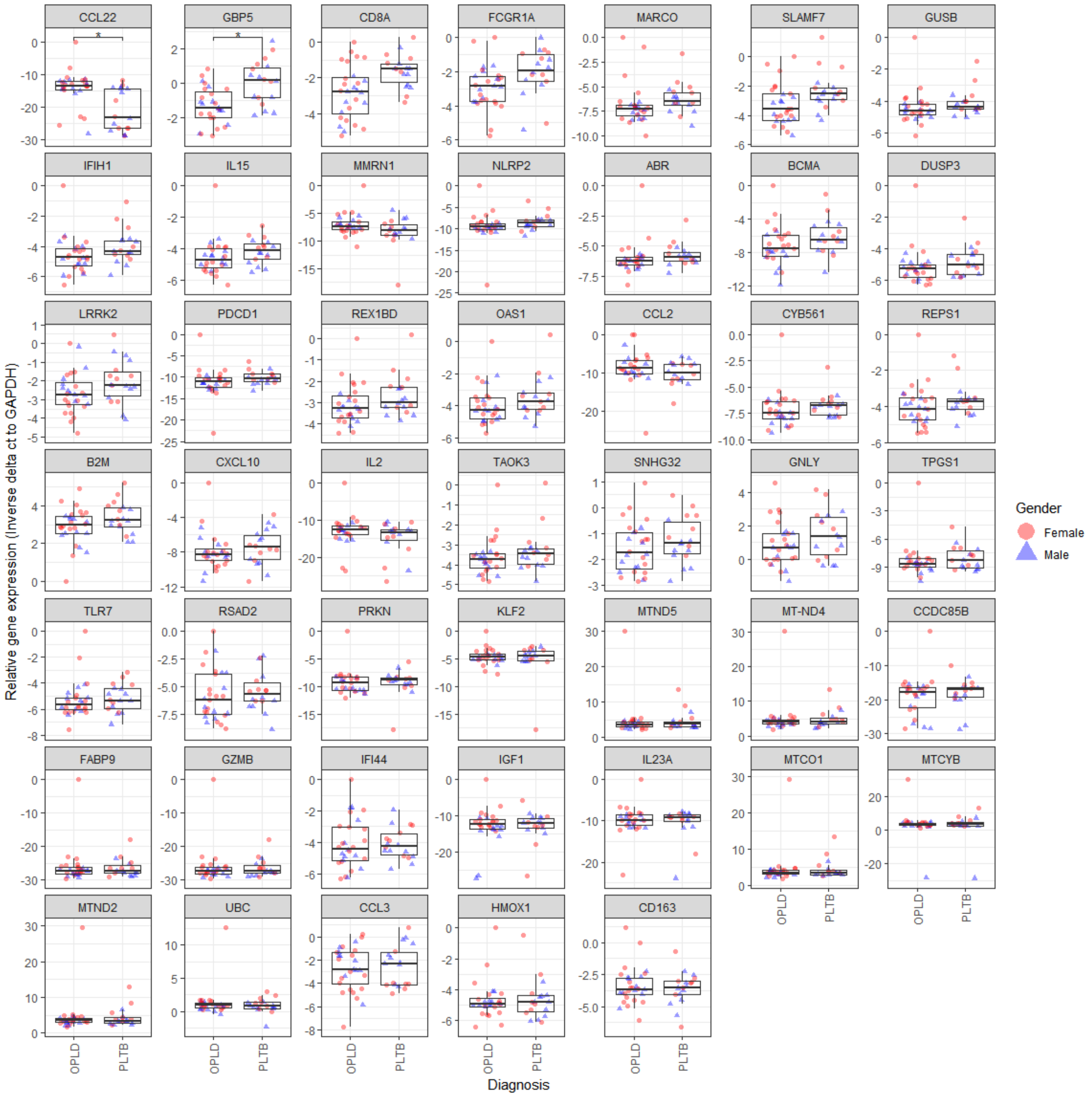


**Supplementary figure 4:** Gene expression in PBMC of patients with PLTB and OPLD. The mRNA levels were measured by microfluidic RNA-seq in human samples of patients with PTLB (n=18), OPLDs (n=28). Relative target gene expression was determined by calculating ΔCt using GAPDH as a reference gene. Since mRNA levels of immune genes were expressed as ΔCt values, larger values represent lower mRNA levels in the sample. For the purpose of obtaining the correct direction, the ΔCt values were multiplied by -1.

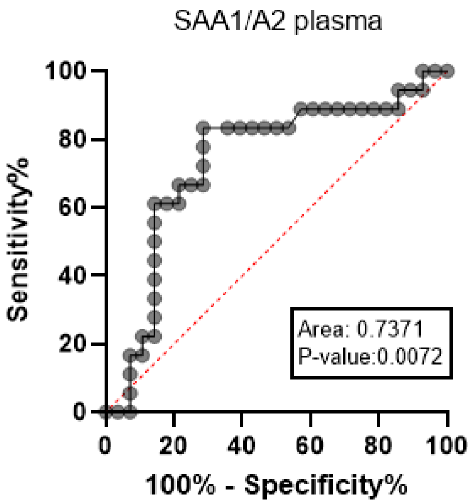


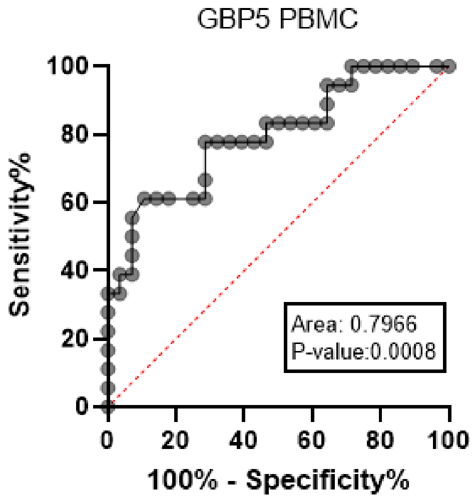


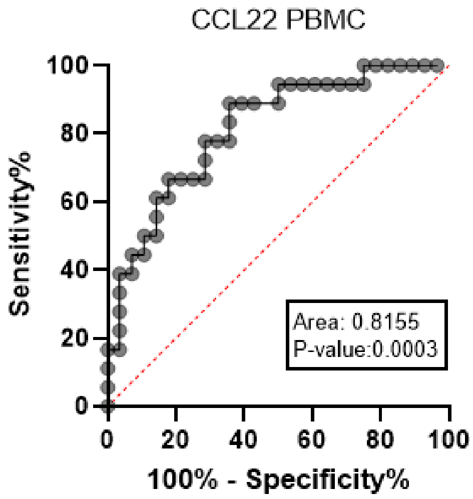


**Blood samples**

| Biomarker | Cut-off | Sensitivity (%) | 95% CI | Specificity (%) | 95% CI |
|---|---|---|---|---|---|
| SAA1A2 | > 1.195 | 83,33 | 60,78% to 94,16% | 71,43 | 52,94% to 84,75% |
| GBP5 | > 0.05000 | 61,11 | 38,62% to 79,69% | 89,29 | 72,80% to 96,29% |
| CCL22 | < -13.83 | 88,89 | 67,20% to 98,03% | 64,29 | 45,83% to 79,29% |

**Blood samples data**

| Biomarker | AUC | p-value |
|---|---|---|
| SAA1A2 | 0,7371 | 0,0072 |
| GBP5 | 0,7966 | 0,0008 |
| CCL22 | 0,8155 | 0,0003 |

**Supplementary figure 5:** Receiver operating characteristic (ROC) curves for the ability of SAA1/A2 (plasma), GBP5 (PBMCs) and CCL22 (PBMCs) to discriminate PLTB from OPLD patients. AUC: area under the curve. Sensitivity and specificity were calculated according to the highest cut-off establish through the Youden's index[1].

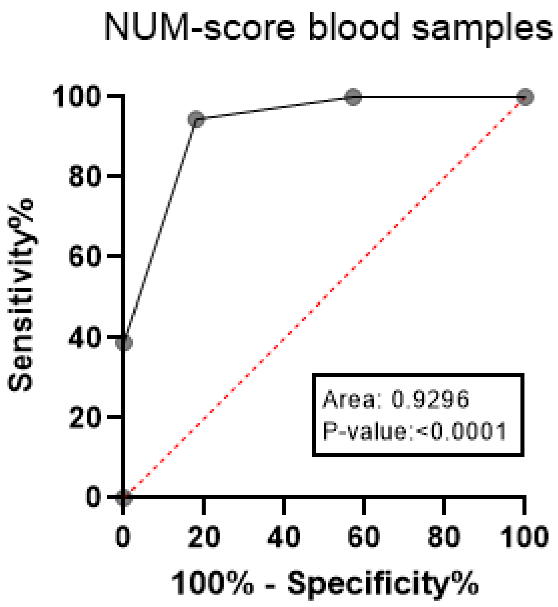


**Blood-derived samples**

| | Cut-off | Sensitivity% | 95% CI | Specificity% | 95% CI |
|---|---|---|---|---|---|
| NUM-score | > 1.500 | 94,44 | 74,24% to 99,72% | 82,14 | 64,41% to 92,12% |

**Blood samples data**

| | AUC | p-value |
|---|---|---|
| NUM-score | 0,9296 | <0,0001 |

**Supplementary figure 6:** Receiver operating characteristic (ROC) curves for the NUM-score of the combination SAA1/A2 (plasma), GBP5 (PBMCs) and CCL22 (PBMCs) to discriminate PLTB from OPLD patients. The NUM-score is based on the sum of the number of individual positive biomarkers detected. This required determination of a cut-off R-value to discriminate PLTB patients from OPLD, which was done using the Youden's index[1] for each individual biomarker. AUC: area under the curve. CI: confidence interval.

**Blood samples**

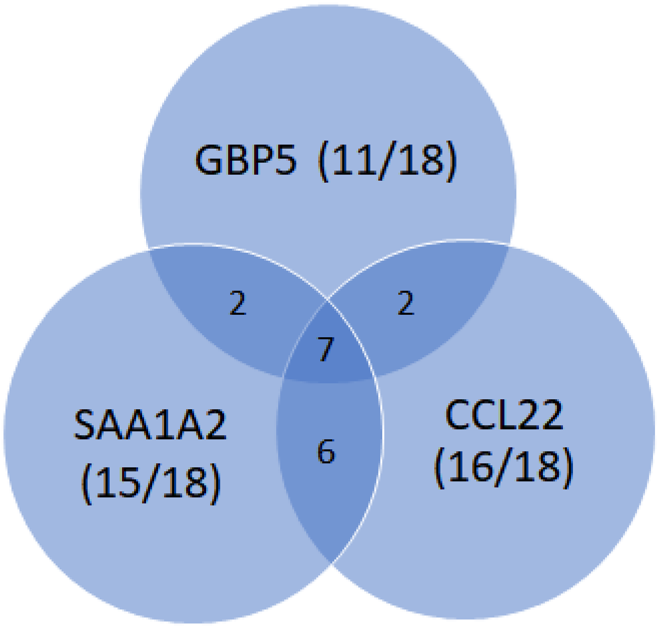


**Supplementary figure 7:** Venn diagram showing the proportion of patients with levels higher than the cut-off level for positivity (based on the Youden index for GPB5, CCL22 and SAA1/A2) in low invasive blood samples. Only individuals of whom both proteomic and transcriptomic markers could be assessed (n=18) were included in the Venn diagram.
